# High Density Phenotypic Map of Natural Variation for Intermediate Phenotypes Associated with Stalk Lodging Resistance in Maize

**DOI:** 10.1101/2025.04.03.647088

**Authors:** Bharath Kunduru, Norbert T. Bokros, Kaitlin Tabaracci, Rohit Kumar, Manwinder S. Brar, Christopher J. Stubbs, Yusuf Oduntan, Joseph DeKold, Rebecca Bishop, Joseph Woomer, Virginia Verges, Armando G. McDonald, Christopher S. McMahan, Seth DeBolt, Daniel J. Robertson, Rajandeep S. Sekhon

## Abstract

The world has food security needs that are currently not being met. Stalk lodging undermines crop productivity and incurs global yield losses of at least $6 billion in maize (*Zea mays* L.). Genetic architecture of stalk lodging resistance, a measure of the ability of the stalk to withstand lodging, remains poorly resolved, creating a bottleneck for genetic improvement. Identification of diverse plant traits at multiple length scales of biological organizations that contribute to stalk lodging resistance and characterization of natural variation for these traits is critical for improving stalk lodging resistance. We identified and evaluated 11 intermediate phenotypes, traits associated with stalk lodging resistance, in a maize diversity panel of 566 inbred lines evaluated over four environments. The identity of each of the 31,260 stalks evaluated in the study was preserved throughout the phenotyping pipeline which enabled capturing variation at the individual plant level. This high-density phenotypic dataset provided a foundation for statistical genomics, predictive modeling, and machine learning analyses to identify genes and genetic elements underlying stalk lodging resistance. Additionally, phenotypic characterization of multiple intermediate phenotypes on a diverse set of inbred lines provided excellent opportunities to understand the relative contribution of these traits to stalk lodging resistance. Besides improvement of maize for grain and animal feedstock, the inferences from this data will be valuable for improvement of stalk lodging resistance in other grass species.

## Context

Stalk lodging, permanent displacement of plant stems from their vertical axis, is a major constraint affecting agricultural crop production and one of the most significant sources of potential calorie losses impacting global food security and economic prosperity. In cereals, stalk lodging can reduce grain yield by 2-80% and significantly deteriorate biomass quality (Kunduru et al., 2023; Lang et al., 2012; Shah et al., 2017). Based on global stalk lodging incidence in maize (FAO, 2008; USDA, 2019), an annual 1% reduction in stalk lodging incidence could feed approximately 5 million more people. Given the substantial economic and food security implications, efforts to enhance stalk lodging resistance, the inherent ability of plant stalks to resist lodging, have focused on both genetic and agronomic approaches. One strategy involves reducing plant height, as short plant stature achieved by altering the gibberellin biosynthesis pathway can decrease the stalk lodging incidence (Stokstad, 2023; Stubbs et al., 2023; Zhao et al., 2022). Dwarf maize hybrids, suitable for high-density planting, are becoming increasingly favored to mitigate stalk lodging and enhance yield levels (Ramstad, 2024). However, high planting densities modify the microenvironment and canopy architecture leading to increased inter-plant competition, higher shading of leaves, and lower photosynthetic potential to paradoxically exacerbate stalk lodging (Sher et al., 2018; Song et al., 2016). Therefore, comprehensive characterization of the genetic architecture of stalk lodging resistance is critical for minimizing yield losses and improving crop productivity in maize.

Stalk lodging resistance is a complex trait controlled by an interplay between several genetic and environmental factors (Bokros et al., 2024; Guo et al., 2021; Xue et al., 2016). The poorly resolved genetic basis of this trait is a serious bottleneck to the improvement of stalk strength in crop plants. A major challenge in genetic studies is the lack of a standardized phenotyping method to consistently and reliably quantify stalk lodging resistance (Kunduru et al., 2023; Tabaracci et al., 2024). Traditionally, field-based phenotypic assessment has relied on late-season lodging incidence determined by counting lodged plants at harvest (Robertson et al., 2016). However, this method is poorly reproducible due to the heavy influence of various environmental factors including wind speed, soil moisture, soil fertility, and pest pressure. Another approach that exposes field-grown plants to high-velocity winds also suffers from major drawbacks including inaccurate phenotypic assessment due to inconsistent wind loading on stems and high cost of construction and maintenance of the phenotyping equipment. To overcome these limitations a novel field-based phenotyping platform, Device for Assessing Resistance to Lodging IN Grains (DARLING), was developed to provide measurements of stalk strength by replicating the mechanical loads and stress patterns experienced by naturally lodged plants (Cook et al., 2016). Using the DARLING, data show that the most reliable and reproducible predictors of stalk lodging resistance of field-grown plants are stalk bending strength and stalk flexural stiffness (Sekhon et al., 2020).

Another major challenge in genetic analysis is that stalk lodging resistance is a complex phenotype that is ultimately determined by structural, material, and geometric properties of stalks manifested at multiple length scales ranging from cell, tissue, organs, and organismal levels (Jiao et al., 2019; Kunduru et al., 2023; Oduntan et al., 2024; Wang et al., 2020). Identification of these intermediate phenotypes could enable precise genetic analyses by mapping key loci and regulatory networks and facilitate targeted breeding strategies to enhance stalk lodging resistance while minimizing trade-offs. However, the identity of these intermediate phenotypes and their relative contribution to stalk lodging resistance remain poorly resolved. We developed and applied several medium-throughput phenotyping approaches to systematically identify and characterize these intermediate phenotypes (Cook et al., 2019; Kunduru et al., 2023; Seegmiller et al., 2020; Tabaracci et al., 2024). Collectively, these advancements lay the groundwork for detailed phenotypic analysis of multiple traits that regulate stalk lodging resistance and genetic analyses to elucidate the underlying regulatory mechanisms.

Here, we present a comprehensive dataset comprised of 11 phenotypes representing multiple length scales collected on a diversity panel of 566 maize inbred lines evaluated across four environments. These inbred lines represent the major heterotic groups of the North American maize germplasm, stiff stalk, non-stiff stalk, iodent, as well as sweetcorn, popcorn, and tropical lines. The inbred lines evaluated in this dataset have been densely genotyped, enabling future analyses to examine genotype-phenotype relationships using contemporary and novel models. This dataset provides detailed characterization of the structural and geometric properties of individual internodes located below the primary ear-bearing node. Importantly, the individual plant identity was preserved throughout the study as opposed to averaging phenotypic data across plots. Traditionally, plot averages of multiple plants are used for phenotypic data, but preserving individual plant identities within each inbred of the diversity panel enhances complex trait mapping by capturing phenotypic variability, increasing statistical power, and enabling the implementation of advanced statistical models. This approach also helps disentangle micro-environmental effects within a plot, leading to more accurate estimates of genotypic effects. Additionally, it facilitates multi-trait correlation analysis and improves covariate control, enhancing the detection of rare variants and gene-environment interactions that averaging might obscure. This dataset offers a valuable resource to the plant community for understanding the natural variation in maize stalk architecture, resolving the genetic regulation of stalk lodging resistance, and informing breeding strategies to improve crop productivity and climate resilience.

## Methods

### Plant material and experimental locations

The present study was conducted on a maize panel consisting of 566 inbred lines, including a subset of the maize Wisconsin diversity panel (Grzybowski et al., 2023; Mazaheri et al., 2019), ex-PVP lines, and other publicly available inbred lines, representing a broad sample of the maize diversity (Table 1, Fig. 1). The panel was grown in 2020 and 2021 at the Clemson University Simpson Small Ruminant Research and Education Center, Pendleton, SC (34°37’26.1"N 82°44’11.1"W) and the University of Kentucky Spindletop farm, Lexington, KY (38°07’43.8"N 84°29’19.2"W) (Supplementary files F1-F4). Weather data for the experimental locations, obtained from the National Climatic Data Center database using Climate Data Online webtool (https://www.ncei.noaa.gov/cdo-web/), indicated diverse environmental conditions at the test locations (Fig. 2, Supplementary file F5). Due to variable germination and quality control measures, the number of inbred lines studied in each environment varied and ranged between 378-504. In each environment, the experiment was planted in a randomized complete block design with two replications and each plot consisting of a single 7.62 m row, 0.76 m row spacing, and a total plot area of 5.8 m^2^. At each location, soil tests were conducted to determine the fertilizer application rates and standard agronomic practices were followed to maintain a healthy crop stand in the experimental plots (Supplementary table T1). Irrigation and pest control measures were applied in a timely manner to ensure optimal plant health while keeping pest and disease pressure under minimum threshold levels.

**Figure 1:**
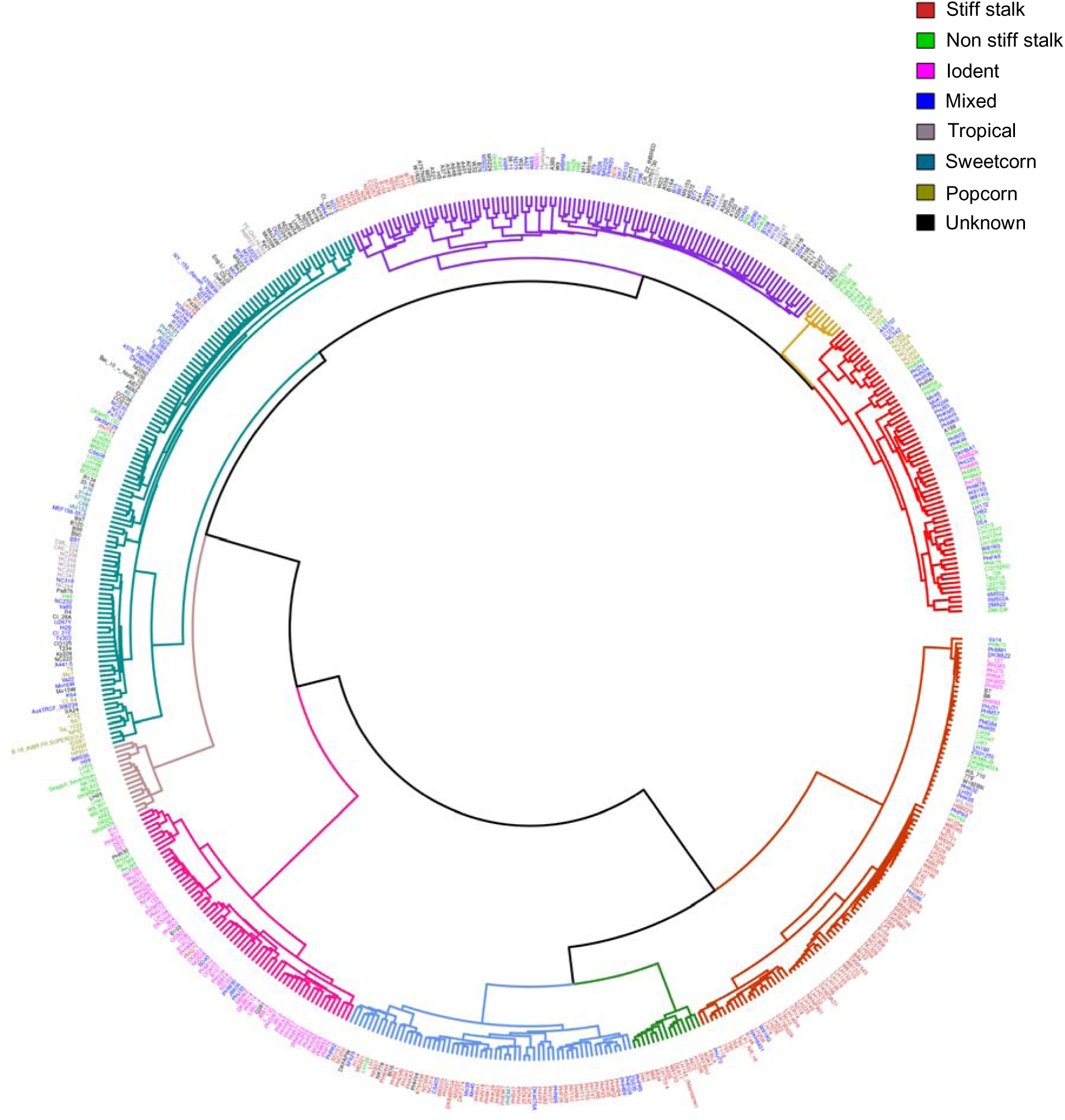
Dendrogram showing phylogenetic relationship among the maize inbred lines evaluated in the present study. The inbred lines representing eight heterotic pools (top right) were classified into nine clusters as indicated by the colored branches. Whole genome resequencing data to construct this dendrogram was obtained from a published study [25]. Only 555 genotypes with available genotype data are shown here.

**Figure 2:**
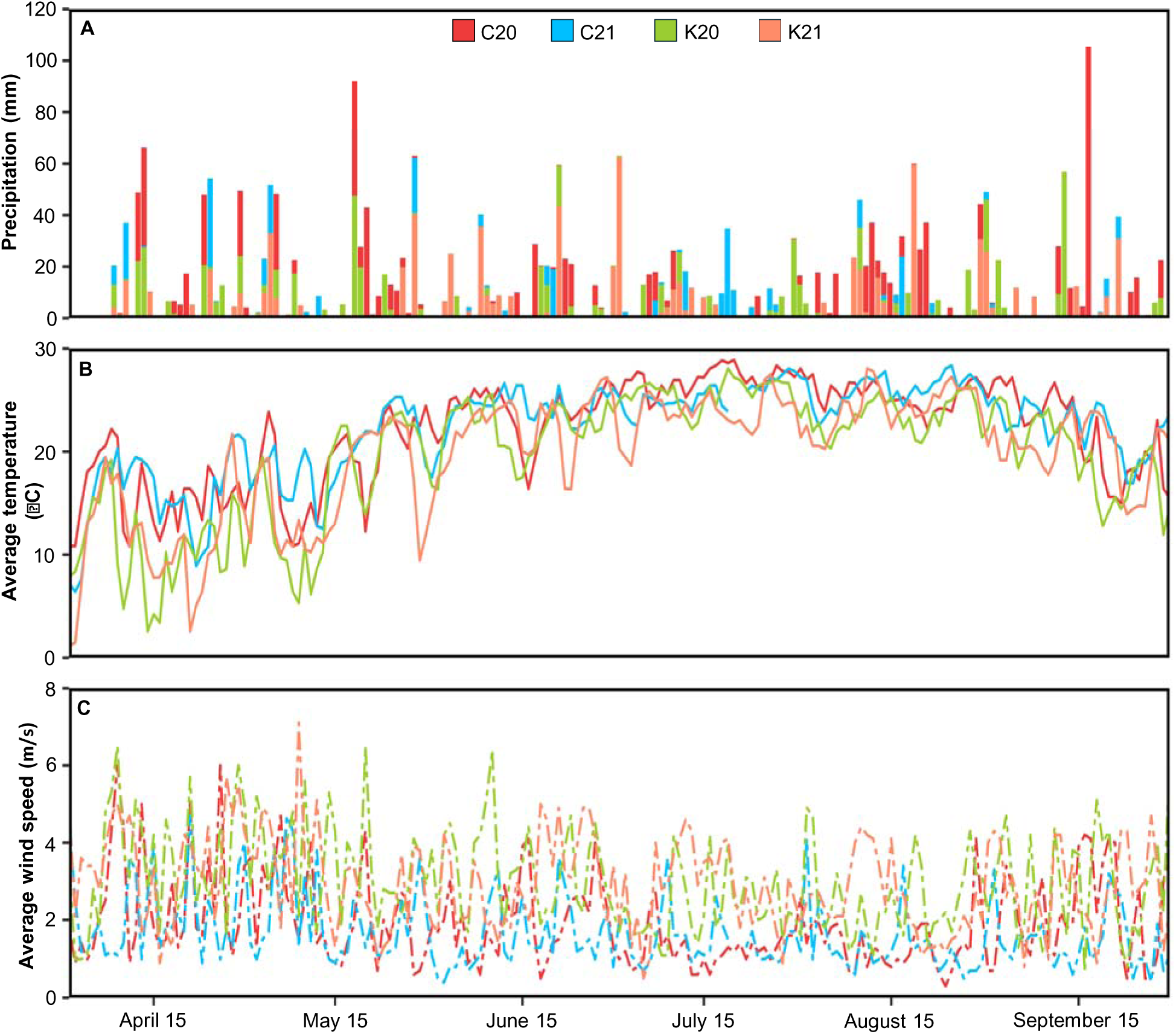
Daily weather distribution at the experimental locations. (A) Total daily precipitation across environments, with color partitioning indicating precipitation amounts at individual locations. (B) Average temperature calculated as the mean of the maximum and minimum temperatures recorded on each date at each environment. (C0 Average wind speed. Abbreviations: C20, Clemson University 2020; C21, Clemson University 2021; K20, University of Kentucky 2020; K21, University of Kentucky 2021; m/s, meters per second.

**Table 1:**
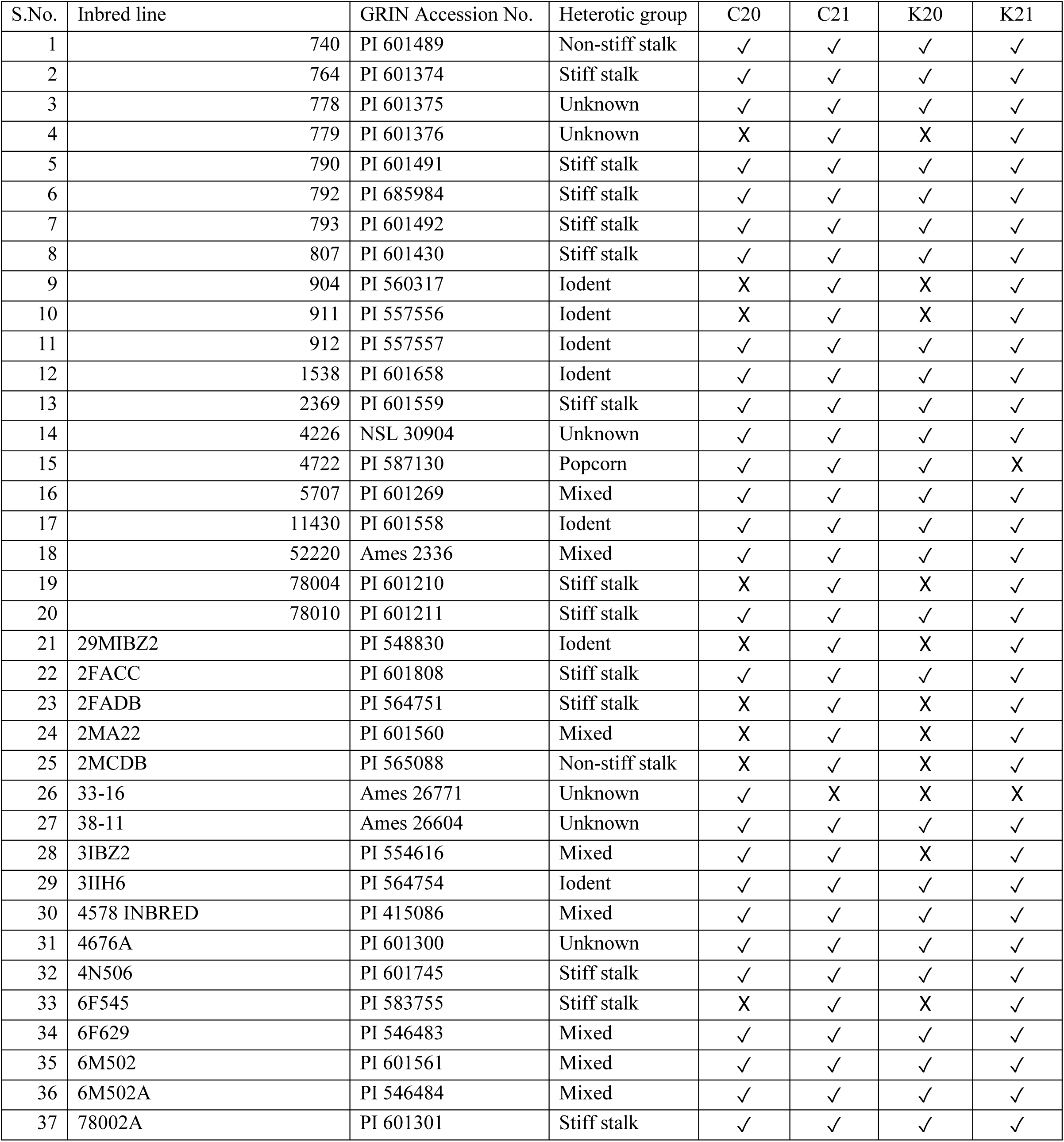
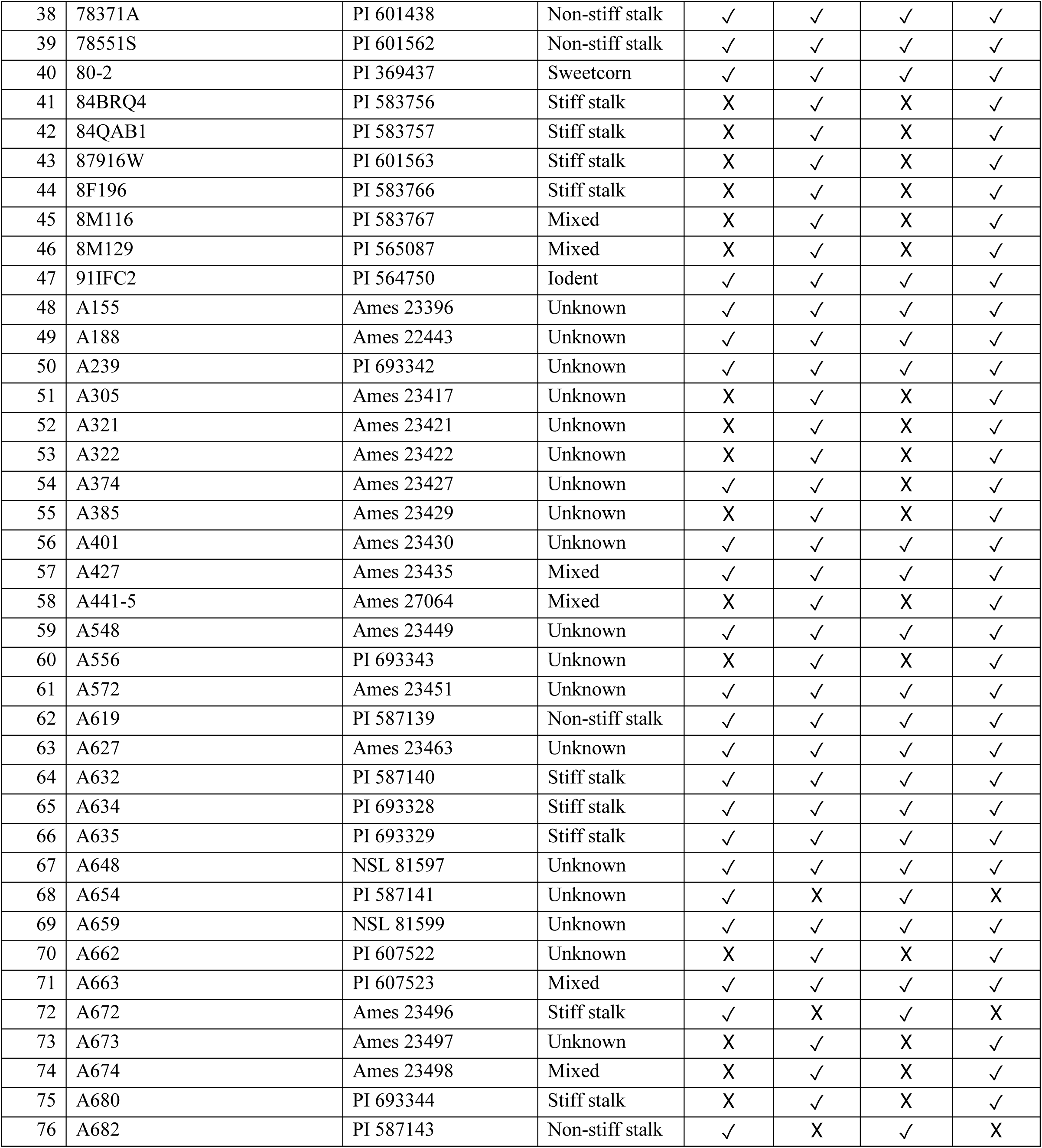
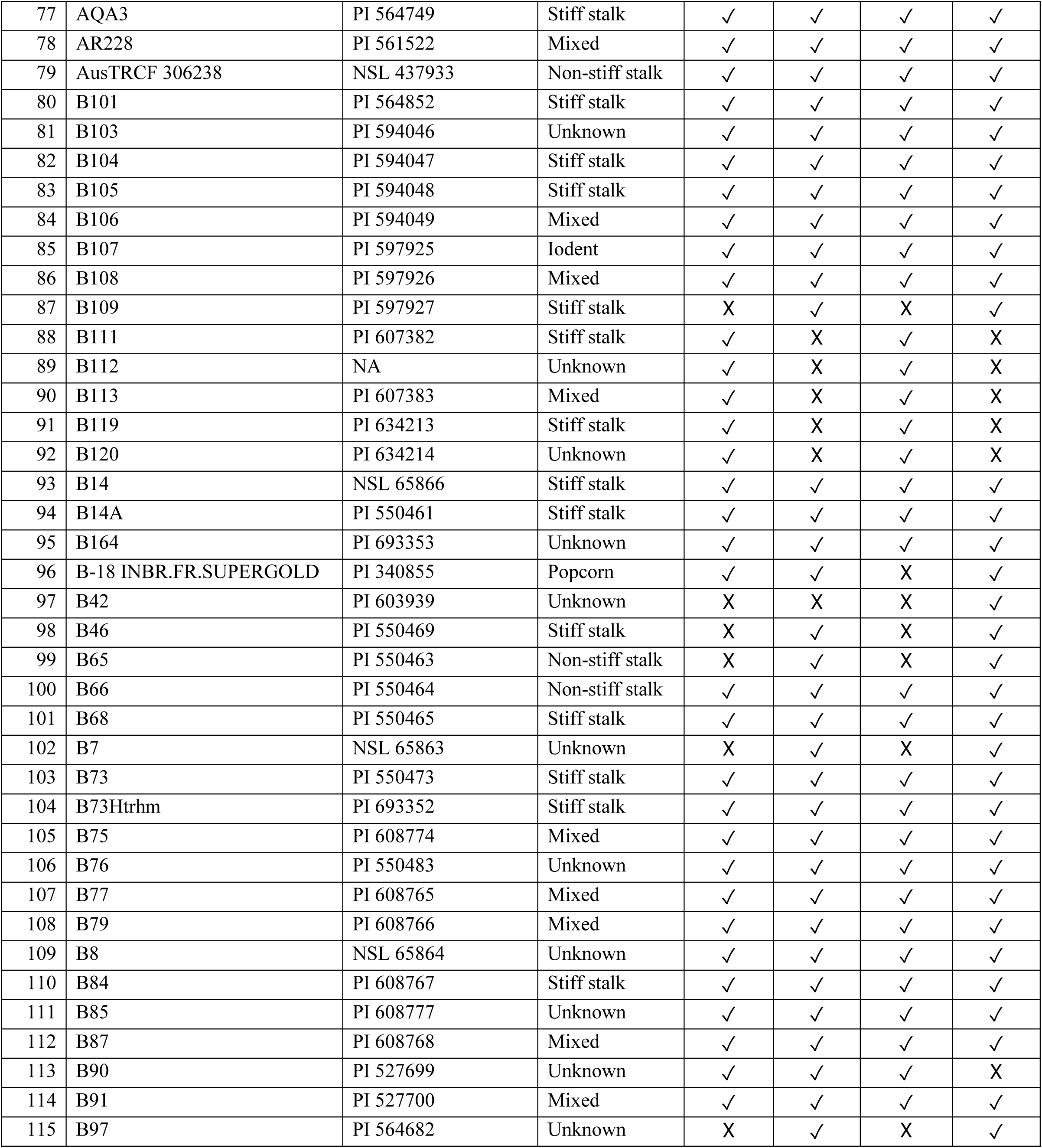
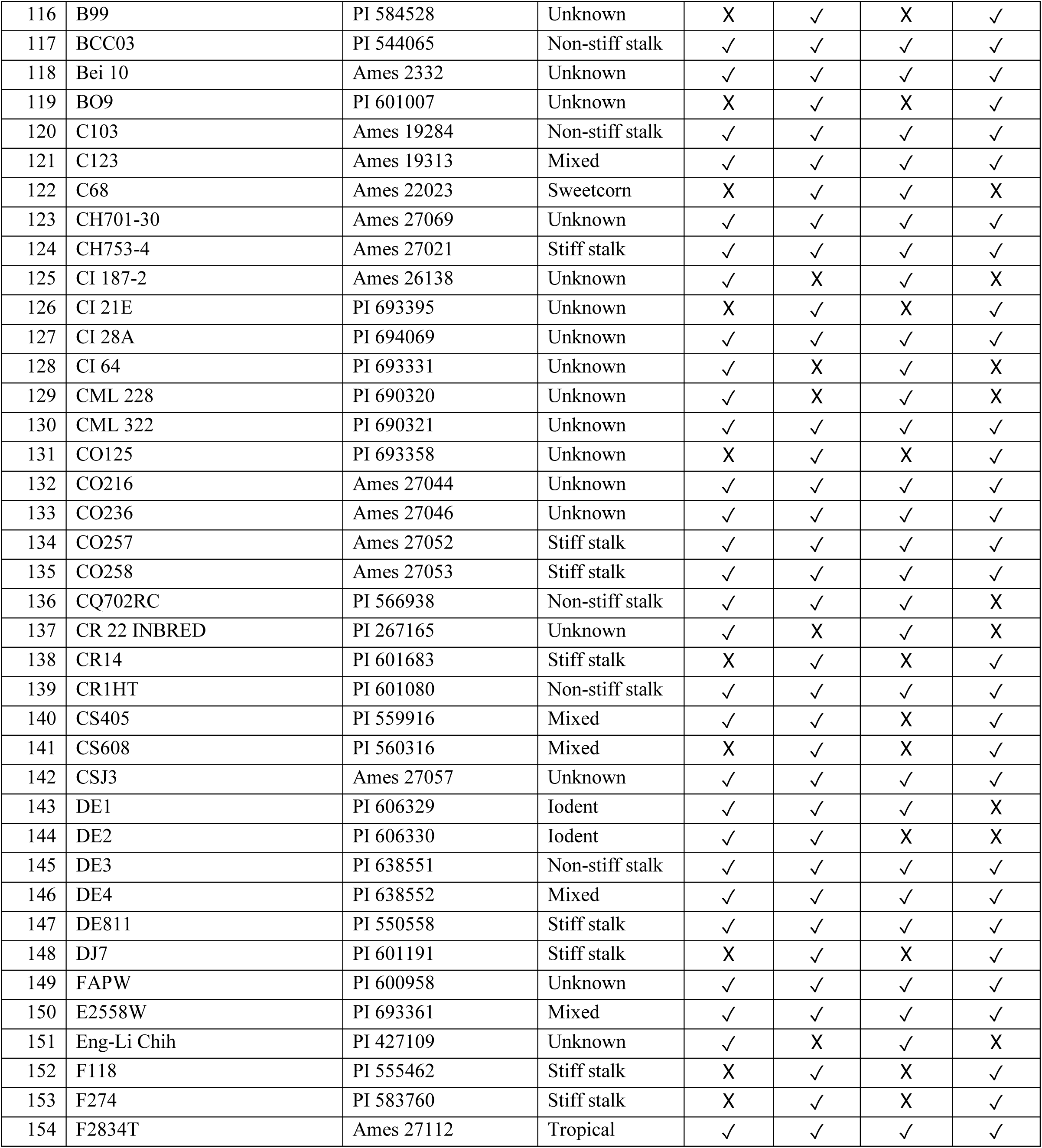
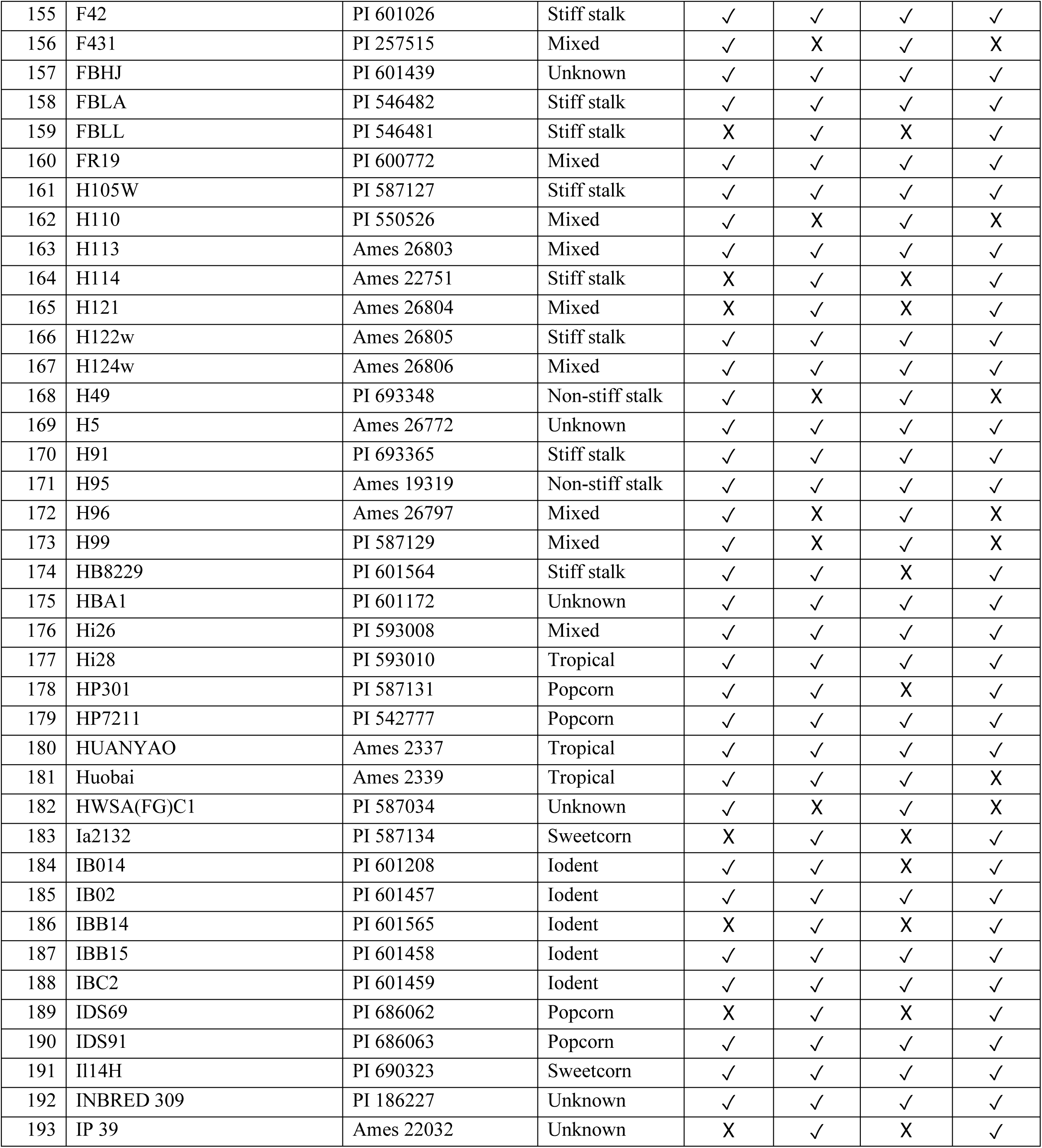
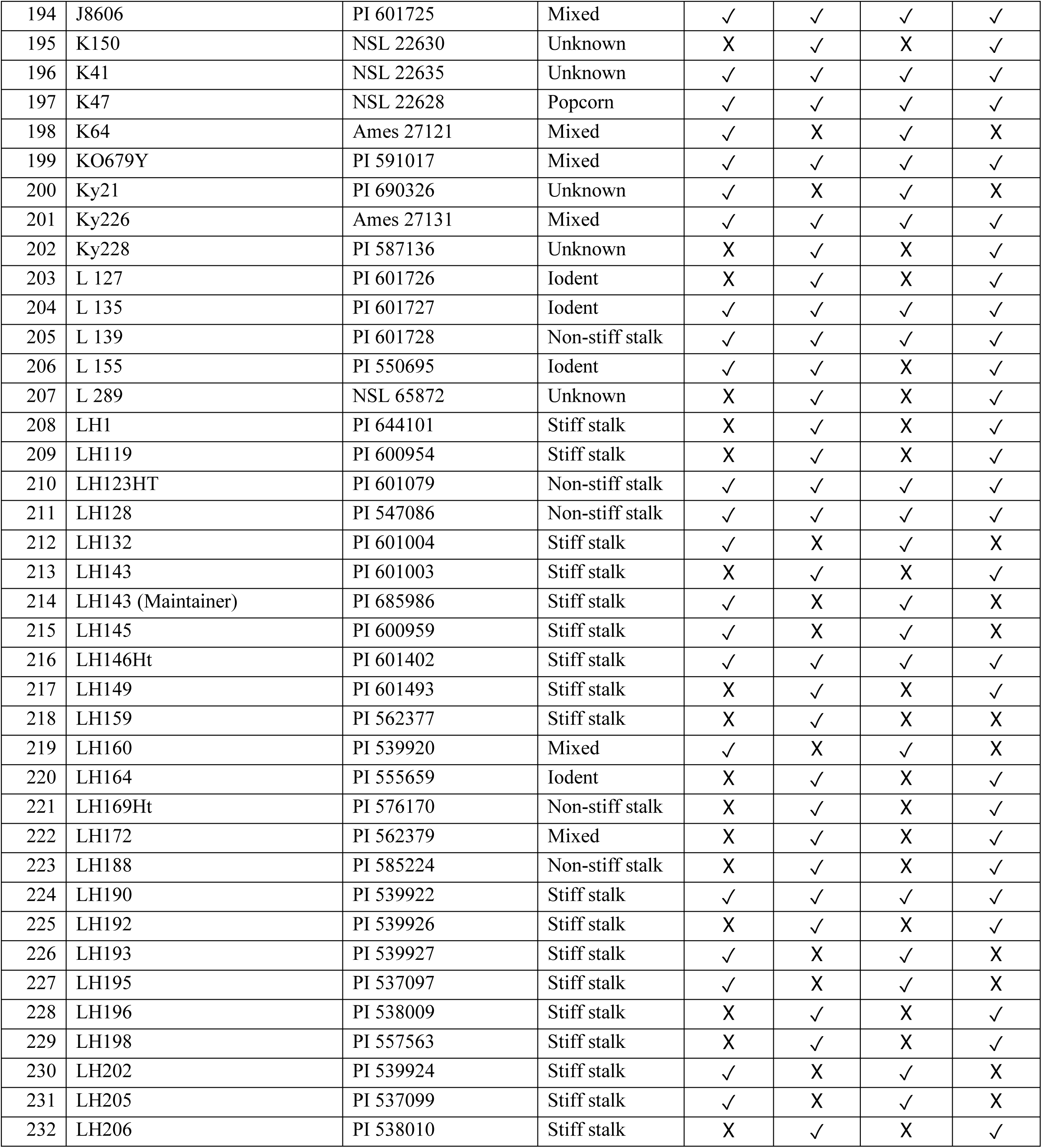
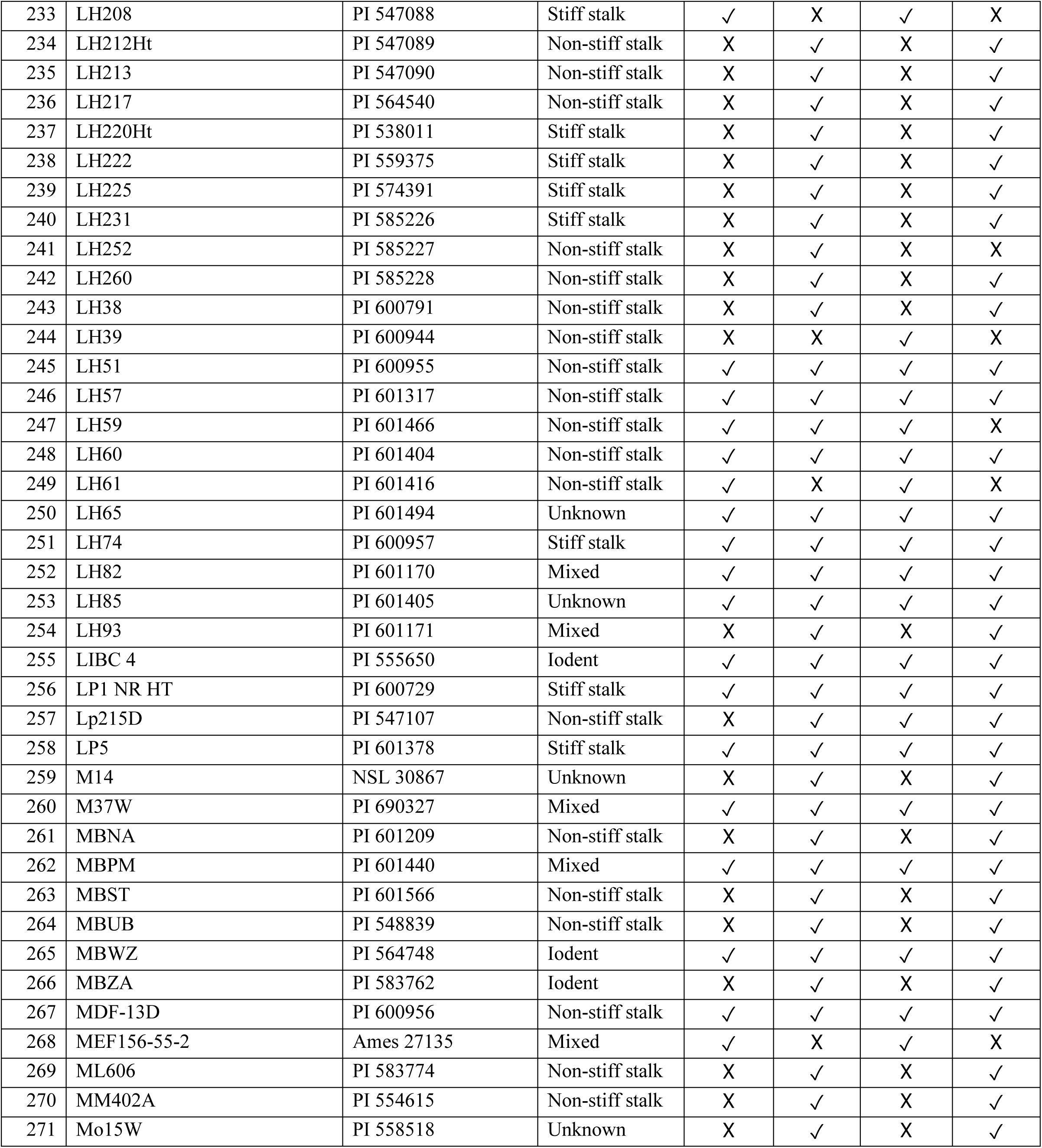
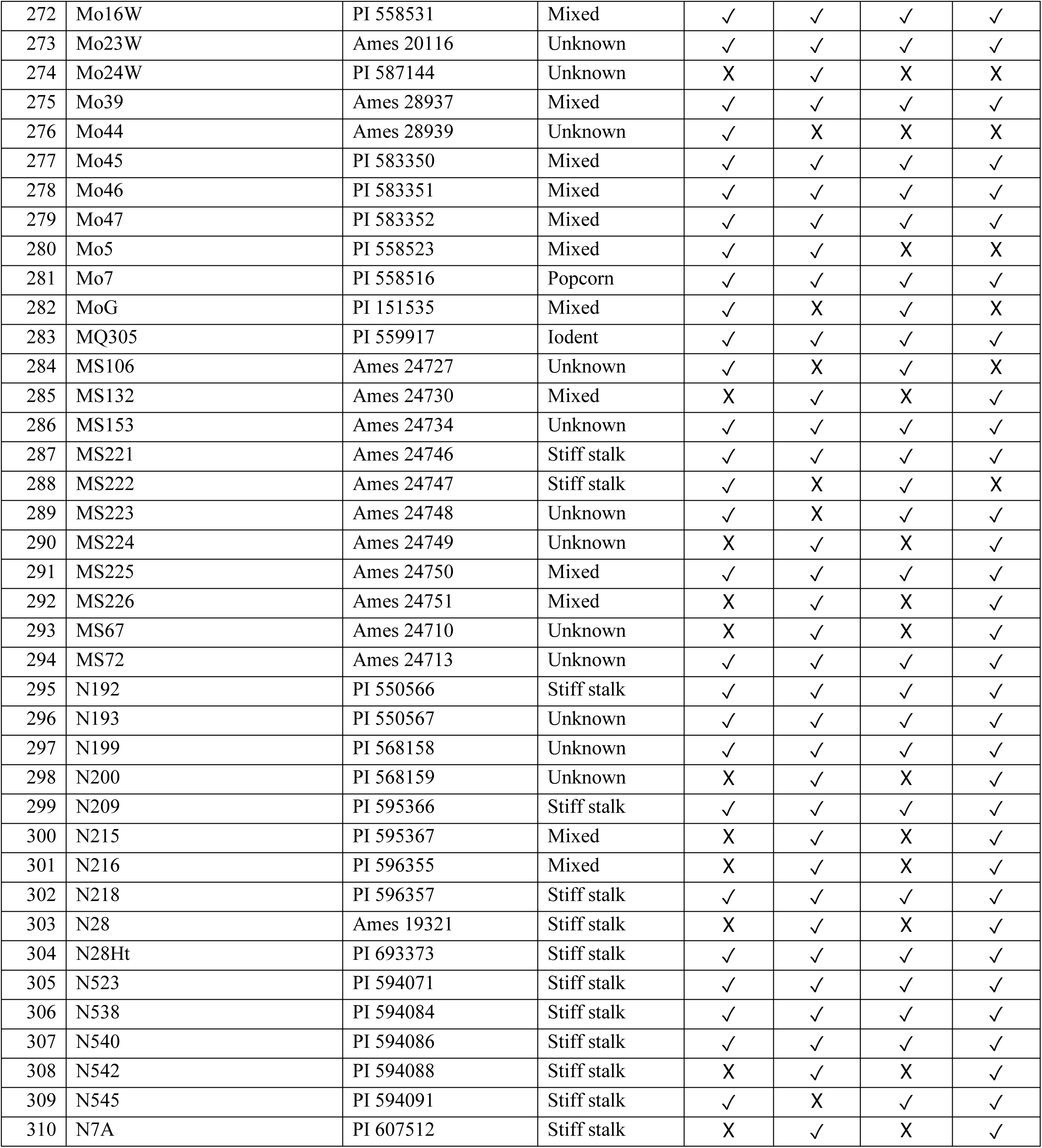
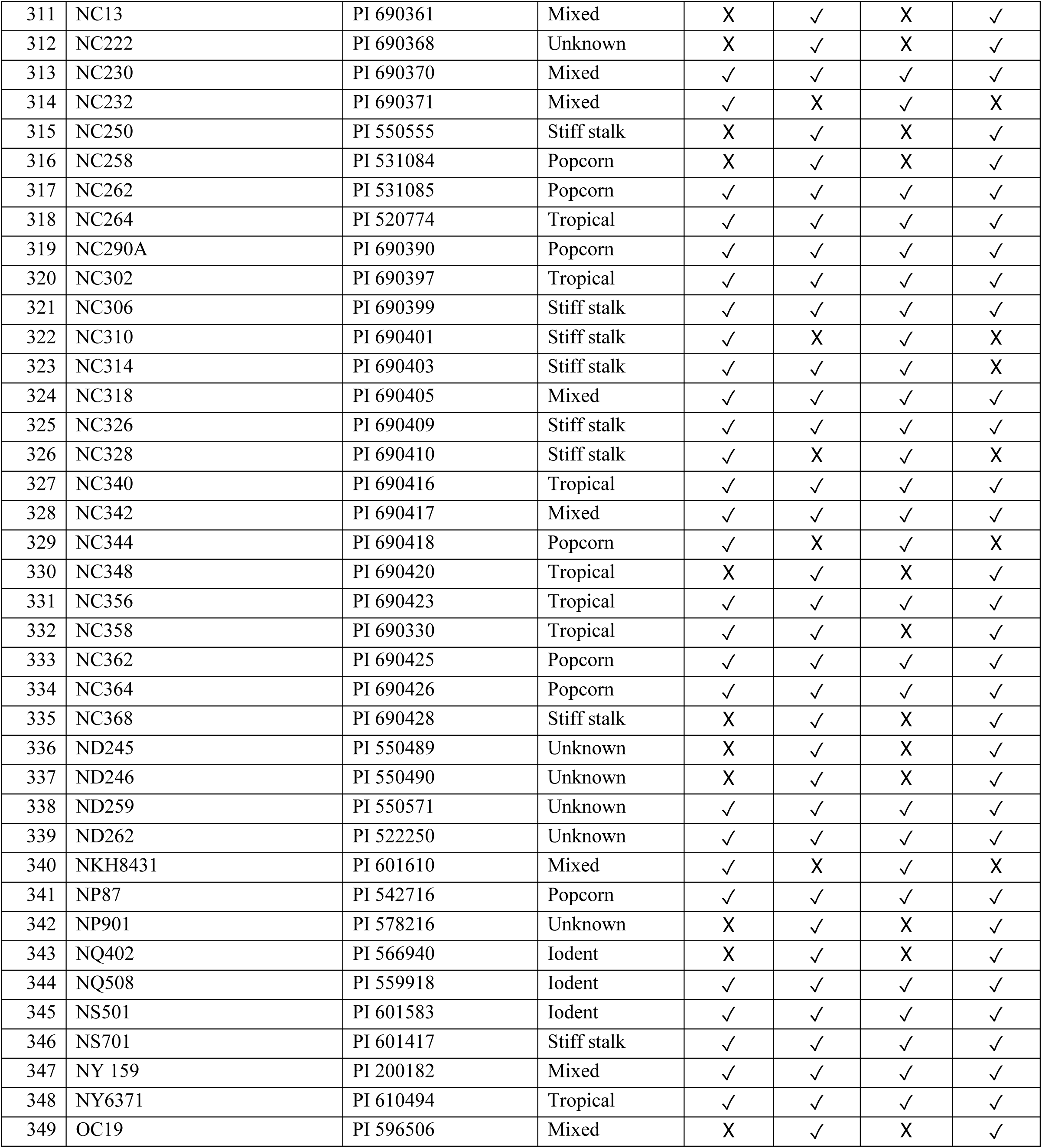
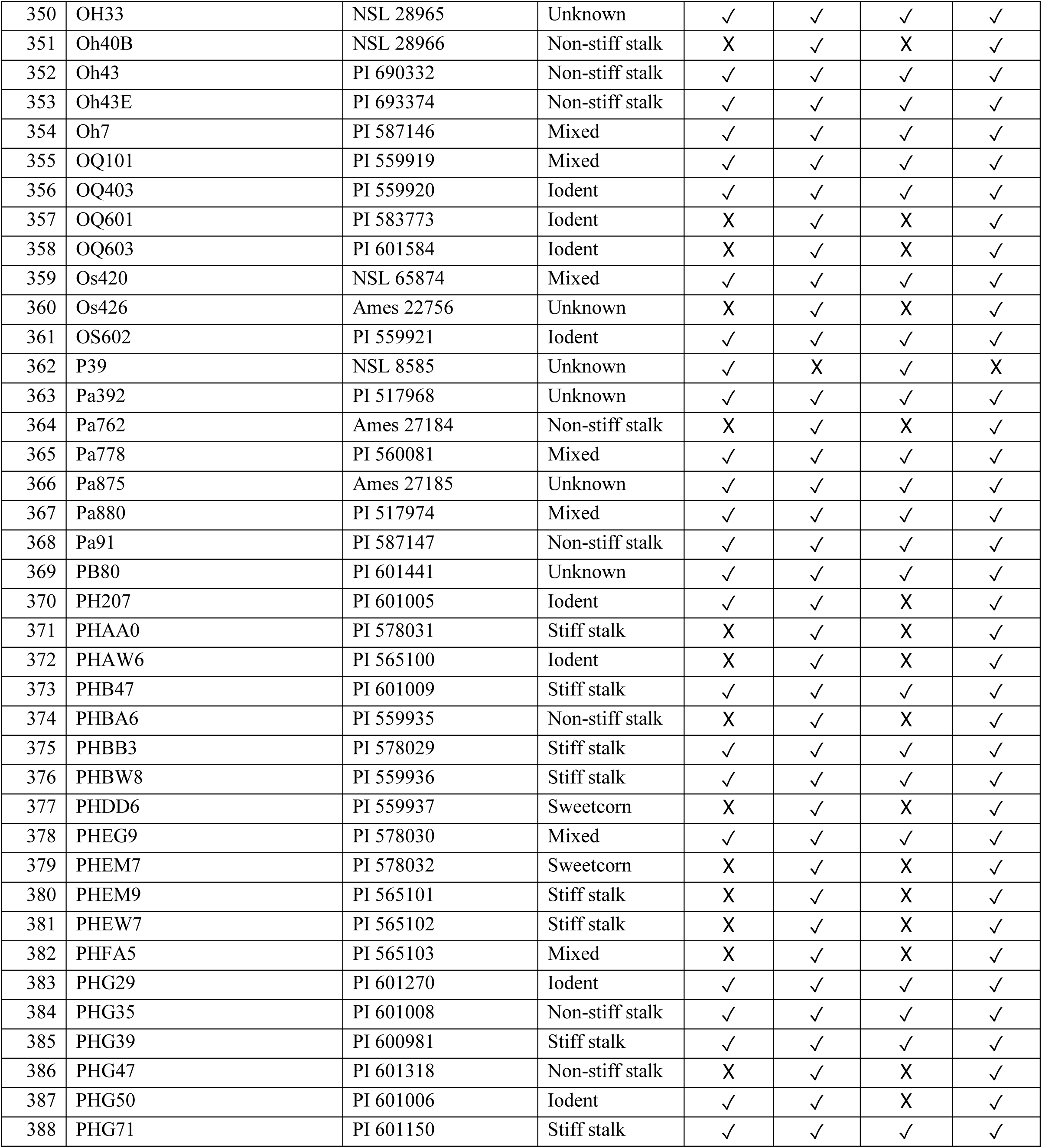
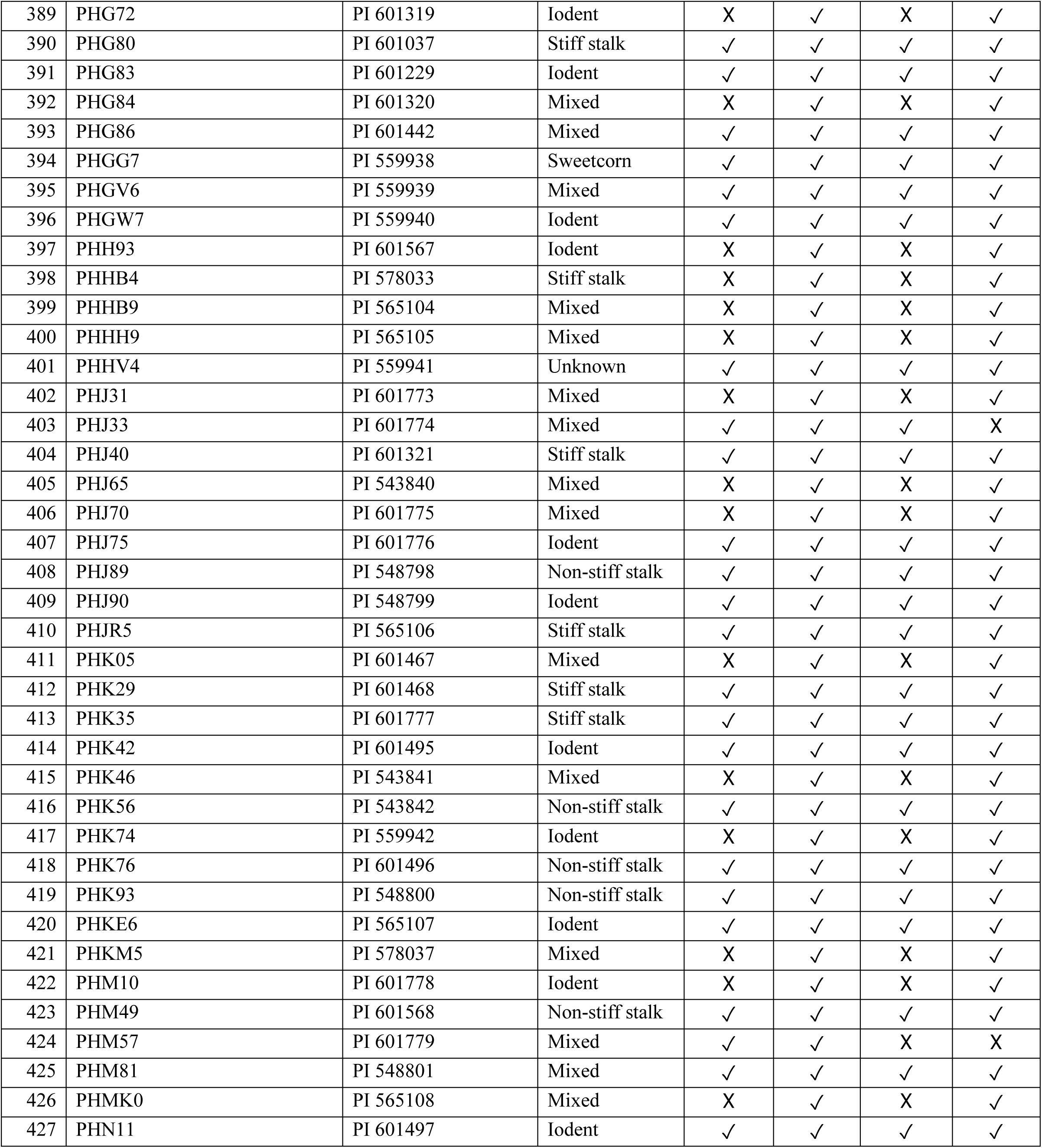
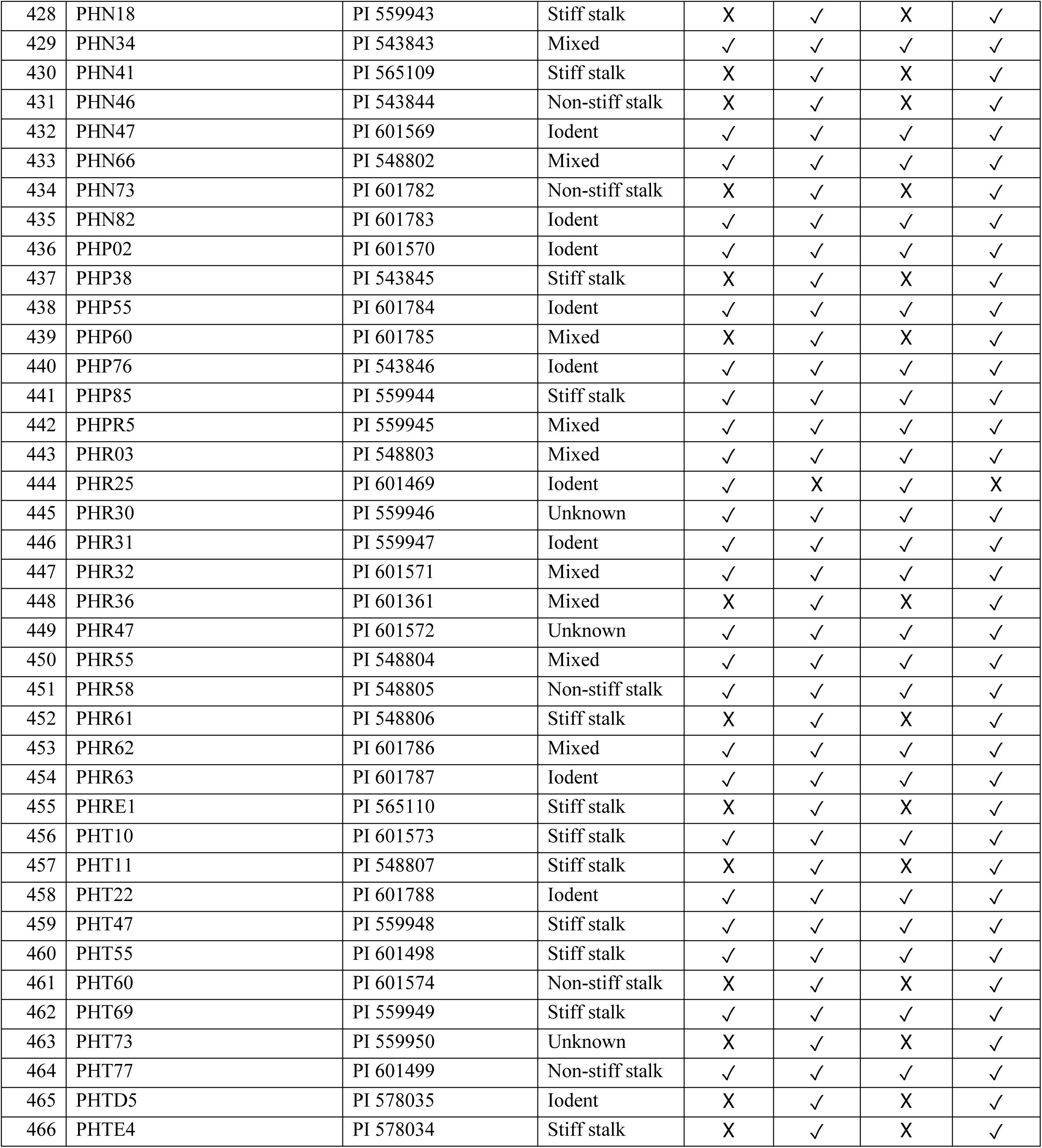
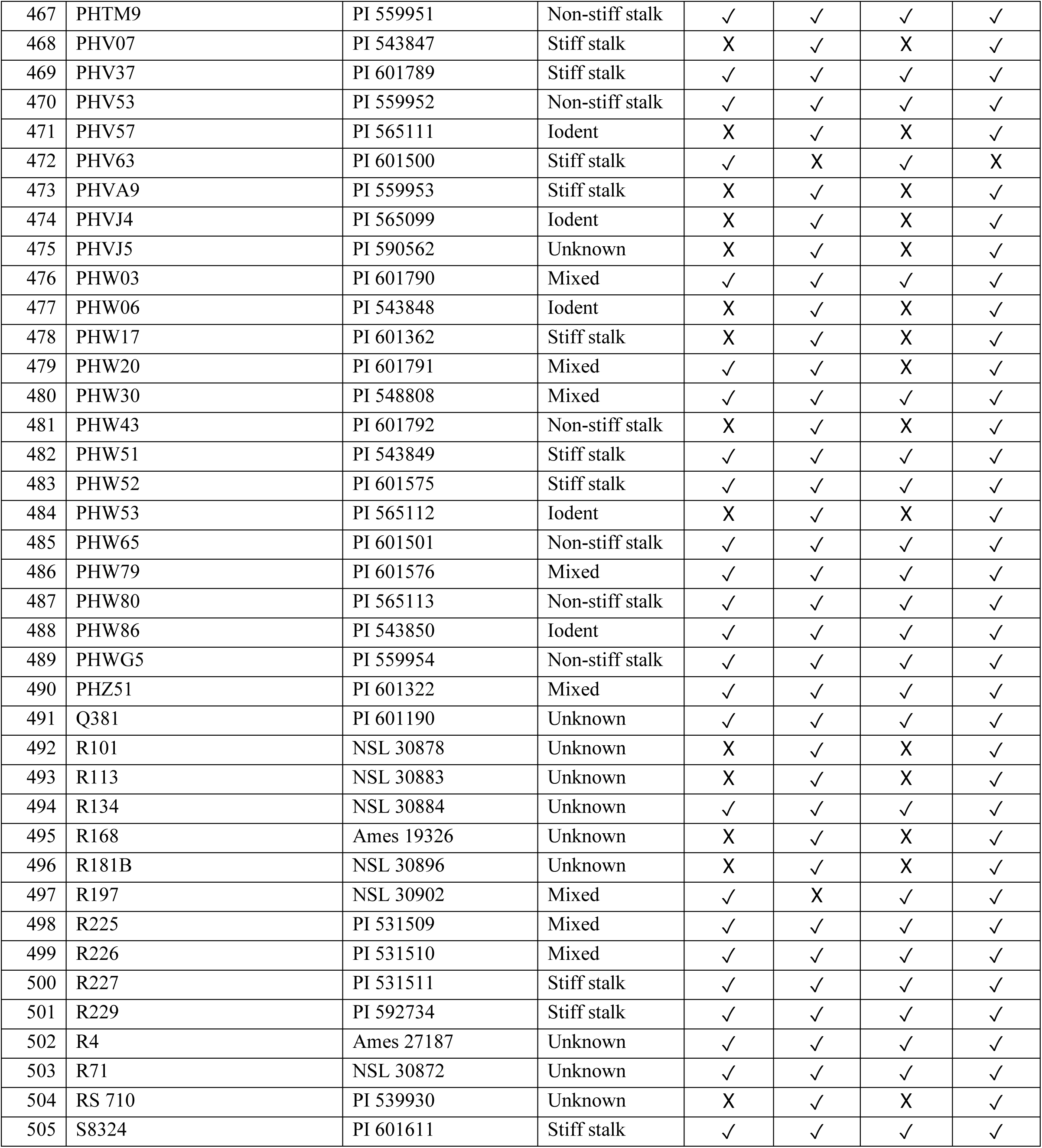
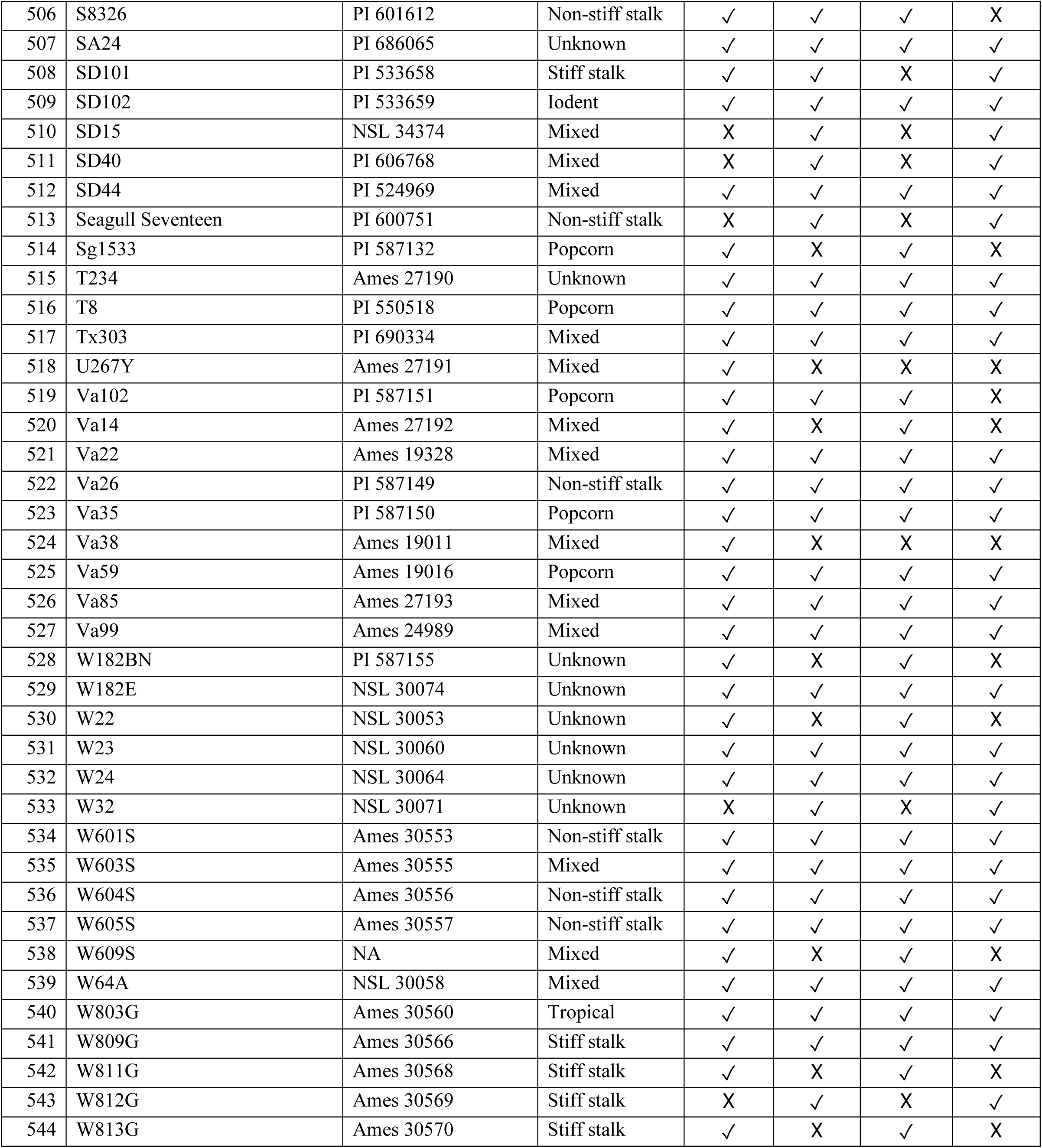
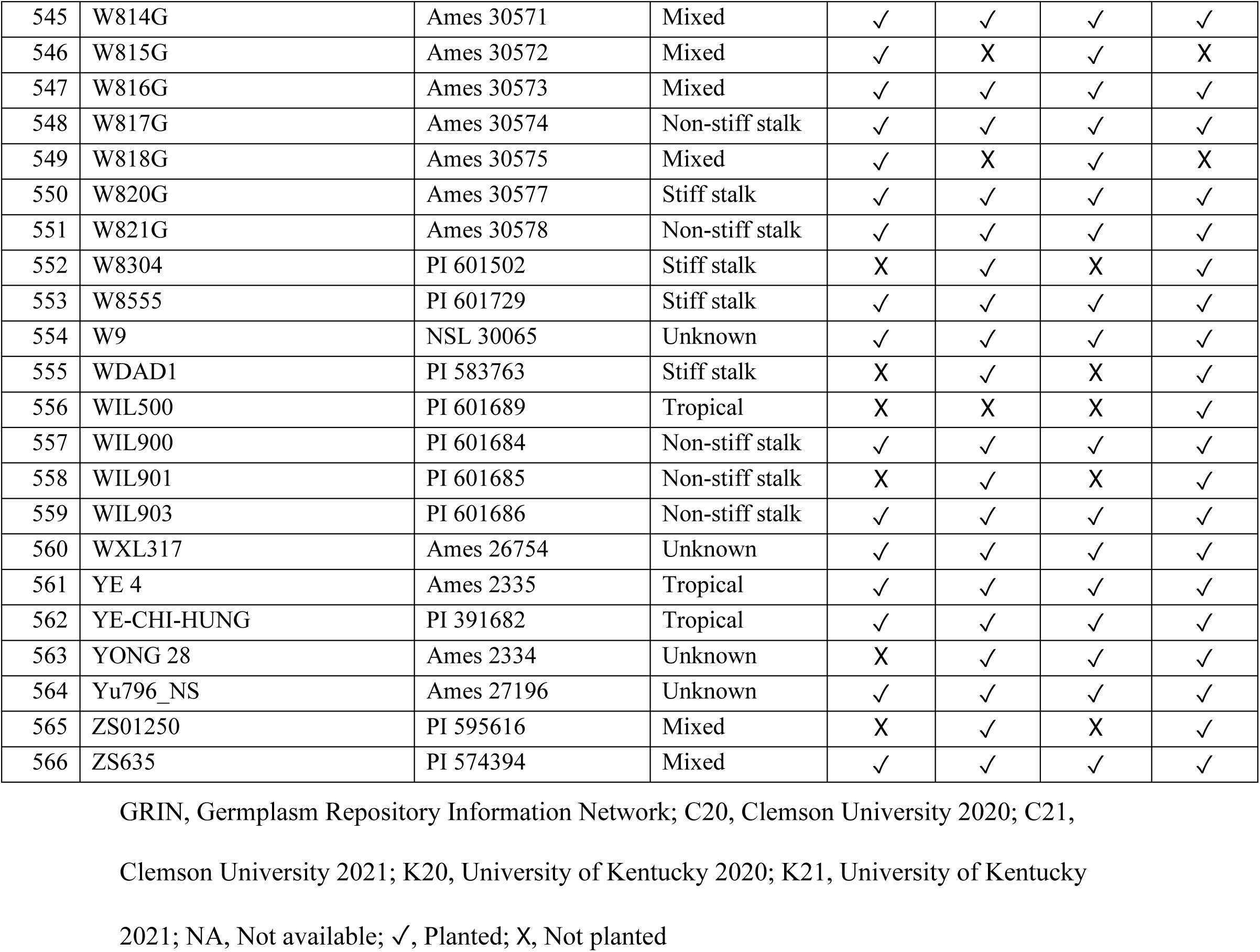
Details of maize inbred lines evaluated in the present study.

### Intermediate phenotypes and sampling strategy

To reach the data we aimed to extract from the experiment the sampling approach needed to consider multiscale measurement with individual plant resolution. Thus, various whole plant- and internode-level phenotypes associated with stalk lodging resistance on individual stalks were recorded while maintaining the stalk identity for each recorded phenotype. Plant-level phenotypes included plant height, ear height, stalk flexural stiffness, and stalk bending strength (Fig. 3). The internode-level phenotypes comprised major diameter, minor diameter, rind thickness, rind penetration resistance, moment of inertia, section modulus, and integrated puncture score experienced by the internodes (Fig. 4-5). Sample preparation, description, and phenotyping protocols for the listed intermediate phenotypes are detailed elsewhere (Kunduru et al., 2023; Stubbs et al., 2022; Tabaracci et al., 2024). Briefly, ten representative and healthy-looking plants per plot were tagged with unique barcode labels to facilitate data matching between field and laboratory measurements. Plant-level phenotypes were collected on live plants in the field, whereas the internode-level phenotypes were measured on the harvested and dried stalks in the laboratory. The phenotyping methodology and relevance of different intermediate phenotypes in assessing stalk lodging resistance are discussed below.

**Figure 3:**
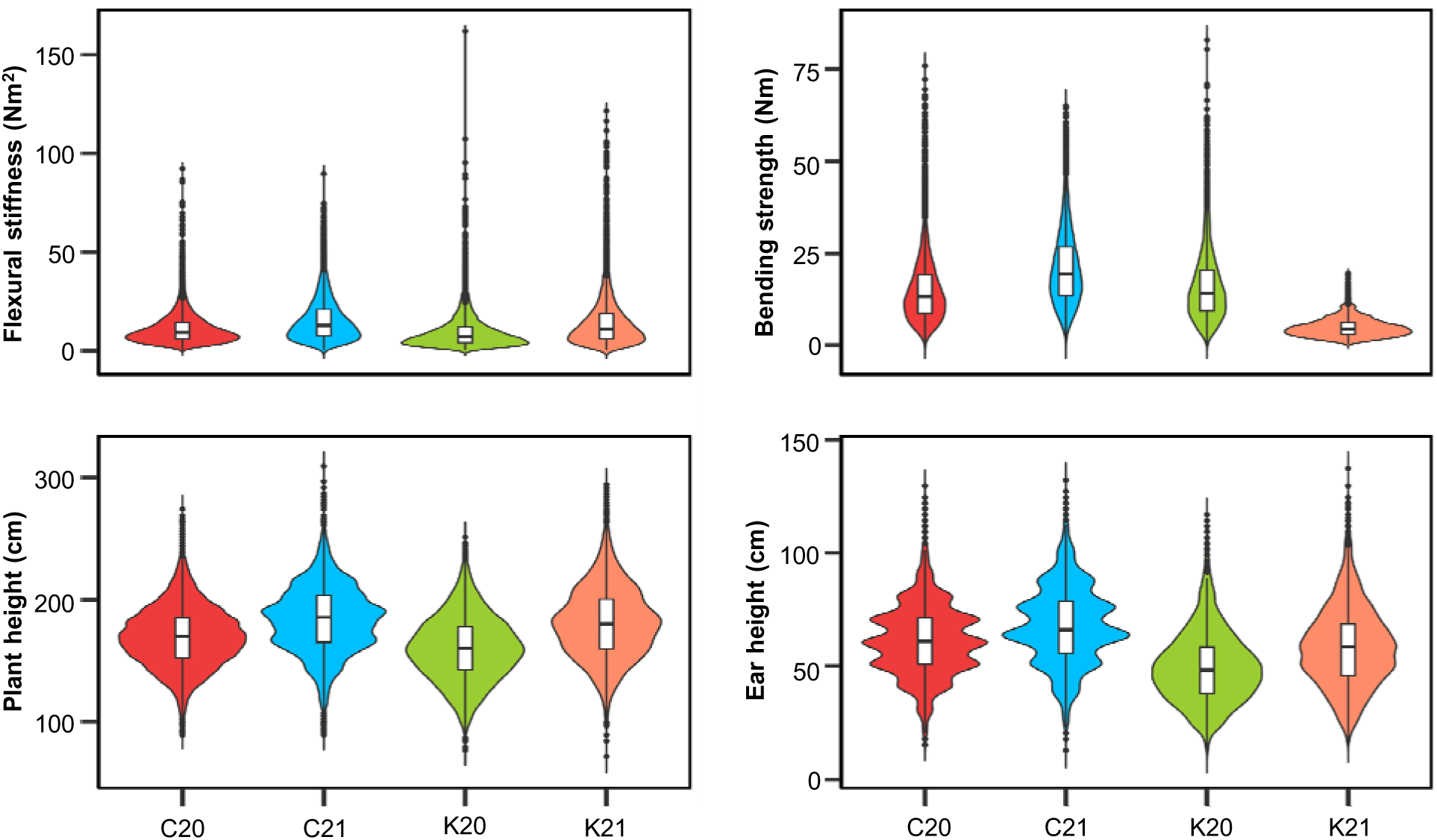
Phenotype distribution of the plant-level intermediate phenotypes associated with stalk lodging resistance. Violin plots represent the distribution of phenotype values in different environments while the embedded box plots illustrate data distribution around the median. Within each boxplot, the horizontal line represents median whereas, lower and upper edges represent the 25^th^ and 75^th^ percentiles, respectively, and the outliers are shown as dots beyond the whiskers. C20, Clemson University 2020; C21, Clemson University 2021; K20, University of Kentucky 2020; K21, University of Kentucky 2021.

**Figure 4:**
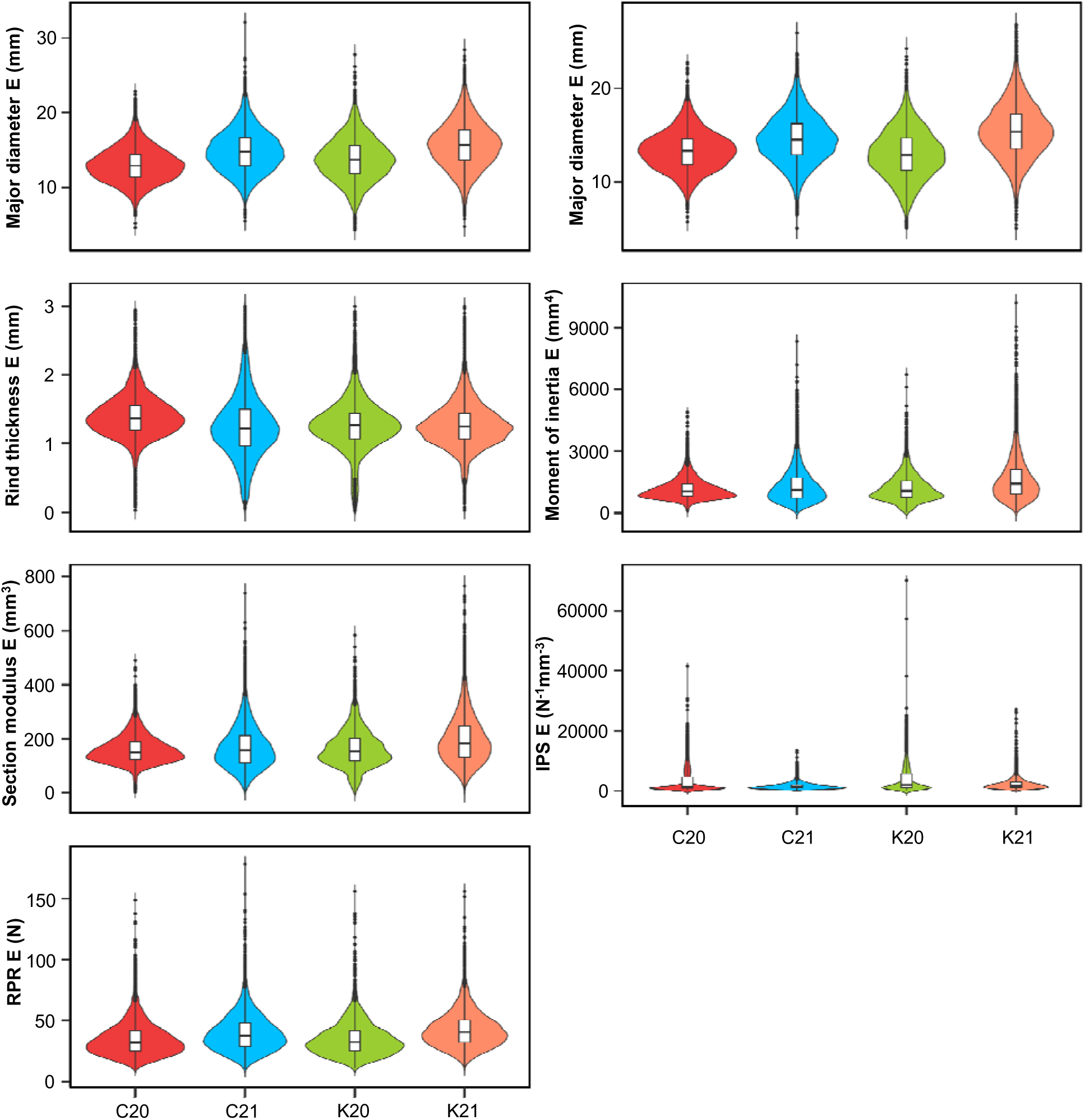
Phenotype distribution of internode-level intermediate phenotypes associated with stalk lodging resistance measured on the internode immediately below the primary ear-bearing node (ear internode). Violin plots represent the distribution of phenotype values in different environments while the embedded box plots illustrate data distribution around the median. Within each boxplot, the horizontal line represents median whereas, lower and upper edges represent the 25^th^ and 75^th^ percentiles, respectively, and the outliers are shown as dots beyond the whiskers. E, Ear internode; IPS, Integrated puncture score; RPR, Rind penetration resistance; C20, Clemson University 2020; C21, Clemson University 2021; K20, University of Kentucky 2020; K21, University of Kentucky 2021.

**Figure 5:**
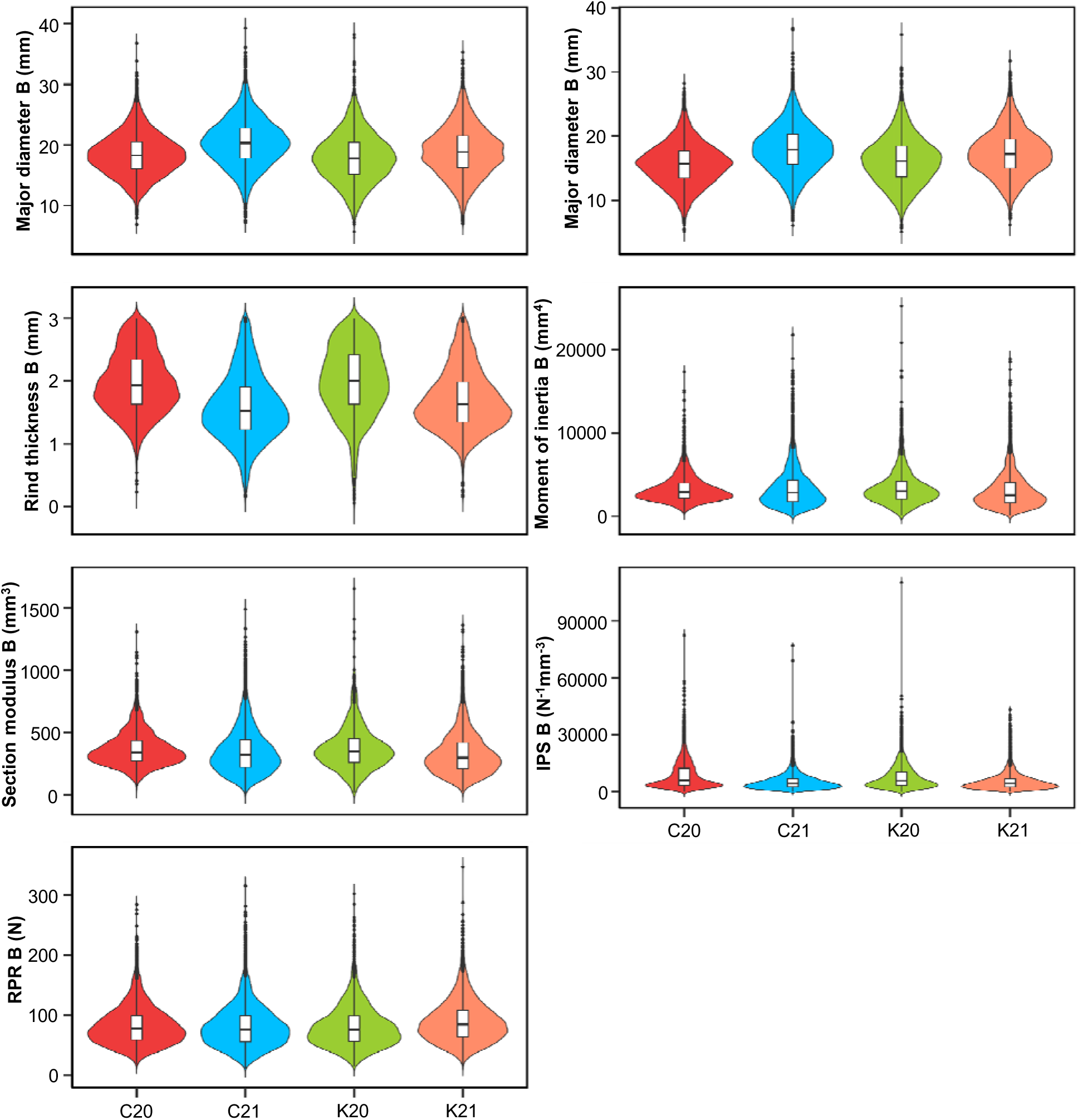
Phenotype distribution of internode-level intermediate phenotypes associated with stalk lodging resistance measured on the first elongated internode above the soil-level (bottom internode). Violin plots represent the distribution of phenotype values in different environments while the embedded box plots illustrate data distribution around the median. Within each boxplot, the horizontal line represents median whereas, lower and upper edges represent the 25^th^ and 75^th^ percentiles, respectively, and the outliers are shown as dots beyond the whiskers. B, Bottom internode; IPS, Integrated puncture score; RPR, Rind penetration resistance; C20, Clemson University 2020; C21, Clemson University 2021; K20, University of Kentucky 2020; K21, University of Kentucky 2021.

### Days to 50% tasseling

In each plot, the number of days from sowing to the initiation of pollen shedding in at least 50% of the plants was recorded.

### Days to 50% silking

In each plot, the number of days from sowing to the initiation of silk emergence in at least 50% of the plants was recorded.

### Plant height

Plant height was recorded one week after days to 50% silking, using PVC pipes marked with metric units, as the linear distance from the tip of the primary rachis of the tassel to the base of the stalk at the ground level. Plant height is a key determinant of stalk lodging resistance with dwarf genotypes being less susceptible to lodging in various cereal crops, including maize (Barten et al., 2022; Gomez et al., 2020; Hirano et al., 2017; Stubbs et al., 2023).

### Ear height

Ear height was recorded as the linear distance from the node bearing the primary ear to the base of the stalk at the ground level. Genotypes with reduced ear height have a lower center of gravity and a shorter lodging arm, both of which enhance stalk lodging resistance (Wen et al., 2019; Zhang et al., 2019).

### Stalk flexural stiffness

Stalk flexural stiffness is a strong predictor of stalk lodging resistance and an effective, non-destructive method for assessing stalk strength under field conditions (Robertson et al., 2016). In the field, stalk flexural stiffness in Nm^2^ was measured 38-42 days after 50% silking with DARLING (Cook et al., 2016; Cook et al., 2019) following established protocols (Kunduru et al., 2023; Tabaracci et al., 2024). For DARLING measurements, the top portions of the plants were removed by excising stalks at the middle of the internode immediately above the primary ear-bearing node. The remaining stalk sections, still rooted in the soil, were prepared by stripping off all lateral appendages to retain bare stalks for measurements. Each prepared stalk was loaded with an incremental force via DARLING until the stalk deflected without breaking or buckling. The reading was discarded if root lodging was observed.

### Stalk bending strength

Stalk bending strength is a reliable predictor of natural stalk lodging incidence, which is used to assess stalk lodging resistance in different crops (Robertson et al., 2022; Sekhon et al., 2020). The methodology for measuring stalk bending strength (a.k.a. maximum bending moment) in Nm was the same as that for stalk flexural stiffness, except that stalks were loaded with incremental force using DARLING until they buckled or broke. Stalk bending strength is a destructive mode for assessing stalk strength and was therefore recorded after measuring stalk flexural stiffness.

### Major diameter

After phenotyping with DARLING, the stalks were cut at the soil-plant interface and air dried in the greenhouse maintained at 38 ⁰C and 35% RH. These dried stalks were used for all the internode-level measurements. Stalk diameter varies around the internode due to the elliptical shape of maize stalks. To account for this variation, two perpendicular diameters, major and minor, were recorded for each internode. Major diameter, measured in mm, refers to the largest possible diameter of an internode measured in the direction perpendicular to the plane of the ear groove (if present) or to the leaf attachment point at the basal node, using a Vernier caliper. Stalk diameter is positively associated with stalk lodging resistance in cereals (Kashiwagi et al., 2008; Zhang et al., 2018).

### Minor diameter

The smallest possible diameter of an internode, measured in mm, was recorded in the direction of the ear groove plane (if present), or the leaf attachment point at the basal node using either a Vernier caliper or an Instron Universal Testing Machine. When measured with the Instron, the probe displacement between the entry and exit points of the stalk tissues was recorded as the minor diameter.

### Rind thickness

In maize stalks, the lack of distinction between rind and pith tissues, which mainly differ for the extent of lignification of parenchyma cell walls (Jung & Casler, 2006), poses challenges for accurately measuring rind thickness. Using the force-displacement curve generated by an Instron Universal Testing Machine during stalk puncture test, we determined rind thickness as the probe displacement from rind-pith transition zone to epidermis (Kunduru et al., 2023; Seegmiller et al., 2020). As the primary mechanical tissue conferring load-bearing capacity to the stalks, thickness of rind plays a crucial role in stalk lodging resistance (Al-Zube et al., 2018; Stubbs, McMahan, et al., 2020).

### Rind penetration resistance

Rind penetration resistance (a.k.a. rind puncture resistance), measured in N, was recorded as the peak force from the force-displacement graph obtained by puncturing the stalk tissues with an Instron Universal Testing System. Rind penetration resistance is a widely used parameter to study and improve stalk lodging resistance in maize (Kumar et al., 2021; Liu et al., 2020; Peiffer et al., 2013).

### Moment of inertia

Moment of inertia, measured in mm^4^, an indicator of the distribution of cross-sectional material in a elliptic cylindrical maize stalk, was calculated from major diameter, minor diameter, and rind thickness using the formula: *I* = *π*[*qp*^3^ − (*q* − 2*r*)(*p* − 2*r*)^3^]/64, where *I* denotes the moment of inertia, *p* and *q* represent major and minor diameter, respectively, and *r* indicates rind thickness. Moment of inertia was reported to be positively associated with stalk lodging resistance and a strong predictor of stalk strength (Kunduru et al., 2023; Robertson et al., 2017).

### Section modulus

Also derived using major diameter, minor diameter, and rind thickness using published protocols and measured in mm^3^. Section modulus had been reported to be a reliable indicator of stalk lodging resistance and not influenced by experimental variables like crop genotype, planting density, etc. (Robertson et al., 2017). Section modulus is equal to the moment of inertia divided by the minor radius of the stalk cross-section.

### Integrated puncture score

Integrated puncture score is a novel force-displacement weighted measurement of rind penetration resistance developed to increase the efficiency of assessing stalk lodging resistance via rind penetration methodologies. This parameter, measured in N^-1^mm^-3^, is derived from the force-displacement graph of stalk puncture tests as per published protocols (Stubbs, McMahan, et al., 2020).

### Naming convention of internodes

The number of elongated internodes on the stalks collected after field phenotyping with DARLING ranged from 4 to 10. To maintain the identity of each internode, we designated these internodes as IN1 to IN10, where IN1 is the internode immediately below the primary ear-bearing node, hereafter referred to as ear internode. Sequentially, the internodes below IN1 were labeled IN2, IN3, and so on, until reaching the bottommost elongated internode, which is labeled as IN*x* and referred to as the bottom internode. For instance, in a stalk with 5 internodes below the primary ear-bearing node, *x* equals 5 and therefore IN5 would be the bottom internode.

### Scripts for data processing

Scripts and example data for calculating rind penetration resistance, internode diameter, rind thickness, and integrated puncture score are available in published literature (Seegmiller et al., 2020; Stubbs, McMahan, et al., 2020). Data analyses, tables, and figures presented in this manuscript were generated using R Statistical Software (version 4.3.1) (Team, 2023).

### Data Validation and quality control Validation

Each phenotyping method utilized to collect data has been validated by separate experiments (Cook et al., 2019; Cook et al., 2020; DeKold & Robertson, 2023; Seegmiller et al., 2020; Stubbs, McMahan, et al., 2020; Stubbs, Oduntan, et al., 2020).

### Quality control

To ensure data uniformity and integrity, we enforced strict quality control measures at every step of phenotyping (Tabaracci et al., 2024). In the field, border plants were excluded from data collection to avoid border effects and mechanically damaged, pest-infested, and/or off-type plants were omitted from phenotyping. After data collection, the entire dataset was compiled into a single database and extensively queried to identify errors. Scatter plots of known phenotypic relationships were generated by year, location, inbred line, and as an aggregated data set to search for outliers or irregular trends. For example, flexural stiffness and bending strength are known to be highly correlated intermediate phenotypes (Robertson et al., 2016), with prior DARLING measurements producing and R^2^ of 0.7 - 0.85. When plotting aggregate flexural stiffness against bending strength, one location produced a significantly different regression slope leading to the discovery of units conversion error. Any outliers were further investigated by reviewing raw data, retaking measurements, or examining the physical stalk specimens for abnormalities. In addition, each phenotype was compared to the ranges observed in prior experiments. For example, rind thickness measurements outside the range of 0.2 to 3.0 mm were flagged for further investigation. The dataset was screened for repeated values, leading to the identification of an instance where the DARLING load cell was maxed out and therefore produced identical values across multiple samples. These quality control measures ensure the accuracy and reliability of the dataset.

### Re-use potential

The stalk phenotype map presented in this dataset is a unique and valuable resource for maize improvement. Collecting such comprehensive data is challenging due to significant cost, labor, and time involved. Generation of this dataset required interdisciplinary expertise to design, implement, and troubleshoot the customized state-of-art phenotyping strategies. Given the logistical complexity and scale of effort, we anticipate that the scientific community will find this dataset valuable. This dataset has broad utility in diverse crop improvement applications, including breeding, genetics, Bayesian and frequentist modeling, machine learning, and software development. This dataset is particularly beneficial to research groups working on maize outside North America who may lack access to the germplasm used in this study. This dataset can be combined with phenotype information from other maize germplasm to further enhance the genetic architecture of maize stalk lodging resistance. The inbred lines included in this dataset are ex-PVP lines or have been in the public domain for several decades. Therefore, the present dataset would be a valuable resource for conducting era studies to compare the dynamics of stalk properties of these historic accessions with recently developed maize germplasm to estimate genetic gains for stalk lodging resistance and related intermediate phenotypes. Moreover, the range of phenotypes included allow researchers to explore the phenotype relationships and trade-offs in breeding maize for bioenergy applications. Beyond biological studies, this dataset also holds engineering relevance for refining stalk geometry and biomechanics, as well as improving phenotyping strategies to enhance stalk lodging resistance in maize.

## Supporting information

Data discussed in manuscript

## Acknowledgements

We sincerely appreciate all undergraduate students and research staff of the Kentucky-Idaho-Clemson Plant Biomechanics Consortium for their support in data collection – William Betsill, Benjamin Clark, Meredith Cobb, Andra Cummings, Bryce Deuty, Grace Gaston, Emma Hatchell, Benjamin Herron, Daniel Hiott, Kaila Honakar, Alexandria Jajack, Rikki Johnson, Grant Kroeschell, Clare Mazzeo, Georgia Moore, Jonathan Tan, Venkata Tatineni and Alison Wuerfel from Clemson University; Lucas Debilius, Anthony DeSantis, Jessy Faulkner, Zane Holiday, Ethan Morris, Nathan LaVoie, Juhyung Lee, Serena Strawn, Nick Locke, Pearl Rwauya, Clayton Bennett, Taylor Spence, Will Seegmiller, Kate Seegmiller, and Andrew Stucker from University of Idaho; Howard Gates, Abigail Haley, Osei Jordan, Jordan Luciano, Nick Rydz, Zoe Schroeder, Ilya Segal, Evanson Telisme, Ren Young, and Jingxia Zhong from University of Kentucky. We also thank the United States Department of Agriculture for providing the germplasm resources used in the study and the farm management crew at Clemson University and University of Kentucky for their assistance in planting and maintaining the research plots. The authors also acknowledge the Clemson University Libraries Open Access Publish Fund awarded to Bharath Kunduru.

## CRediT authorship contribution statement

**Bharath Kunduru –** Methodology, Investigation, Data curation, Formal Analysis, Software, Visualization, Validation, Writing – original draft, Writing – review & editing; **Norbert T. Bokros –** Methodology, Investigation, Data curation, Formal Analysis, Software, Validation, Writing – review & editing; **Kaitlin Tabaracci –** Methodology, Investigation, Data curation, Formal analysis, Software, Validation, Writing – original draft, Writing – review & editing; **Rohit Kumar –** Methodology, Investigation, Writing – review & editing; **Manwinder S. Brar –** Methodology, Investigation, Writing – review & editing; **Christopher J. Stubbs –** Methodology, Investigation, Formal analysis, Writing – review & editing; **Yusuf Oduntan –** Methodology, Investigation, Formal analysis, Writing – review & editing; **Joseph DeKold –** Methodology, Investigation, Formal analysis; **Rebecca Bishop –** Investigation, Writing – review & editing; **Joseph Woomer –** Investigation; **Virginia Verges –** Investigation; **Armando G. McDonald –** Methodology, Funding acquisition, Project administration, Writing – review & editing; **Christopher S. McMahan –** Conceptualization, Data curation, Formal Analysis, Investigation, Methodology, Visualization, Validation, Supervision, Funding acquisition, Project administration, Writing – original draft, Writing – review & editing; **Seth DeBolt –** Conceptualization, Methodology, Investigation, Formal analysis, Supervision, Resources, Funding acquisition, Project administration, Writing – review & editing; **Daniel J. Robertson –** Conceptualization, Methodology, Investigation, Data curation, Formal analysis, Supervision, Resources, Funding acquisition, Project administration, Writing – original draft, Writing – review & editing; **Rajandeep S. Sekhon –** Conceptualization, Methodology, Investigation, Data curation, Formal analysis, Supervision, Resources, Funding acquisition, Project administration, Writing – original draft, Writing – review & editing

## Data availability

All the data associated with this study are available in the Supplementary material.

## Disclaimer

Any opinions, findings, and conclusions or recommendations expressed in this material are those of the author(s) and do not necessarily reflect the views of the National Science Foundation. All the germplasm used in the study are publicly available and we followed the local, national, and international guidelines.

## Abbreviations

⁰C: degrees Celsius
cm: Centimeter
ex-PVP: Expired Plant Variety Protection Act certificate
KY: Kentucky
mm: Millimeter
N: Newton
Nm: Newton*meter
PVC: Polyvinyl chloride
RH: Relative humidity
SC: South Carolina

